# A comparison of GABA-ergic (propofol) and non-GABA-ergic (dexmedetomidine) sedation on visual and motor cortical oscillations, using magnetoencephalography

**DOI:** 10.1101/2020.03.10.985242

**Authors:** Neeraj Saxena, Alexander D. Shaw, Lewys Richmond, Adele Babic, Krish D. Singh, Judith E. Hall, Richard G. Wise, Suresh D. Muthukumaraswamy

**Affiliations:** Cardiff University Brain Research Imaging Centre (CUBRIC), School of Psychology, Cardiff University, Cardiff CF24 4HQ, UK; Department of Anaesthetics, Intensive Care and Pain Medicine, Cwm Taf Morgannwg University Health Board, Llantrisant CF72 8XR, UK; Department of Anaesthetics, Morriston Hospital, Swansea, SA6 6NL, UK; Department of Anaesthetics, Royal Gwent Hospital, Newport, NP20 2UB, UK; Department of Anaesthetics, Intensive Care and Pain Medicine, School of Medicine, Cardiff University, Cardiff CF14 4XW, UK; Institute for Advanced Biomedical Technologies, Department of Neuroscience, Imaging and Clinical Sciences, “G. D’Annunzio University” of Chieti-Pescara, 66100, Chieti, Italy; School of Pharmacy, Faculty of Medical and Health Sciences, Auckland University, Auckland 1123, New Zealand; School of Psychology, Faculty of Medical and Health Sciences, Auckland University, Auckland 1123, New Zealand

**Keywords:** Magnetoencephalography, brain waves, neurophysiology, propofol, dexmedetomidine, conscious sedation

## Abstract

**Background:** Understanding the effects of anaesthetic drugs on cortical oscillations can help to elucidate the mechanistic link between receptor physiology and their clinical effects. Propofol produces divergent effects on visual cortical activity: increasing induced gamma-band responses (GBR) while decreasing stimulus-onset-evoked responses) ^1^. Dexmedetomidine, an α2-adrenergic agonist, differs from GABA-ergic sedatives both mechanistically and clinically as it allows easy arousability from deeper sedation with less cognitive side-effects. Here we use magnetoencephalography (MEG) to characterize and compare the effects of GABAergic (propofol) and non-GABA-ergic (dexmedetomidine) sedation, on visual and motor cortical oscillations.

**Methods:** Sixteen male participants received target-controlled infusions of propofol and dexmedetomidine, producing mild-sedation, in a placebo-controlled, cross-over study. MEG data was collected during a combined visual and motor task.

**Results:** The key findings were that propofol significantly enhanced visual stimulus induced GBR (44% increase in amplitude) while dexmedetomidine decreased it (40%). Propofol also decreased the amplitudes of the M100 (27%) and M150 (52%) evoked responses, whilst dexmedetomidine had no effect on these. During the motor task, neither drug had any significant effect on motor GBR or movement related beta de-synchronisation (MRBD). However, dexmedetomidine increased (92%) post-movement beta synchronisation/rebound (PMBR) power while propofol reduced it (70%).

**Conclusions:** Dexmedetomidine and propofol, at equi-sedative doses, have contrasting effects on visual stimulus induced GBR, visual evoked responses and PMBR. These findings provide a mechanistic link between the known receptor physiology of these sedative drugs and their known clinical effects and may be used to explore mechanisms of other anaesthetic drugs on human consciousness.

## Introduction

Our understanding of the mechanisms of anaesthesia and the neural correlates of anaesthesia-induced unconsciousness is incomplete. A range of theories of anaesthetic mechanism point towards a breakdown of communication between key brain regions as a common endpoint in anaesthesia related unconsciousness ^2^. Oscillatory synchronisation in different frequency bands contributes to long-range neural communication. Of these oscillations, those in the high frequency band (gamma band (30-80 Hz)) are considered key for information processing in the brain ^3^. Studying changes in these neural oscillations provides an opportunity to explore the systems-level mechanistic underpinnings of anaesthetic drug effects and to link them with their known receptor-level effects.

Traditionally, electroencephalography (EEG) has been used to study the human brain’s electrical activity, in relation to anaesthetic mechanisms and its effects on consciousness due to its widespread availability, lower cost and relative ease of use. Magnetoencephalography (MEG) is a neuroimaging technique where the changes in the magnetic field induced by the electrical activity in the brain are recorded. The external magnetic field is generated by slow postsynaptic potentials in the dendrites of the cerebral cortex. Similar to EEG, the dendritic processes need to be spatially aligned in order to generate measurable fields, making the apical dendrites of pyramidal cells the most likely sources of MEG/EEG activity. The main advantage of MEG over EEG is that the neuromagnetic signal is unaffected by the difficult to measure conductivity profile of the skull and scalp making it more robust in localising current source generators. Coupled to the fact that commercial MEG can use hundreds of channels, its spatial resolution is significantly greater than a typical 32/64 channel EEG set-up, although high density EEG systems are increasingly being used (for more in-depth review on MEG and pharmaco-MEG see Baillet ^4^ and Muthukumaraswamy ^5^, respectively). Finally, and importantly in the context of the current study, MEG is significantly more sensitive than EEG in detecting high-frequency gamma-band activity and better at separating gamma-band activity from the muscle artefacts that can contaminate this band ^6^.

While most of the research into anaesthetic mechanisms using neuroimaging have focused on temporal oscillatory activity during rest, task-related oscillatory changes may also provide further mechanistic insights into the actions of these drugs. Sustained narrow band gamma band oscillations are generated in the visual cortex in response to a simple visual contrast pattern. These patterns arise from the interactions between the excitatory and inhibitory neural networks, which shape both the amplitude and peak frequency of these gamma oscillations. According to the pyramidal-interneuron gamma (PING) model, the local interaction of superficial pyramidal cells and inhibitory interneuron populations underlies oscillations in the gamma-frequency band (30+ Hz) (see Buzsaki and Wang ^3^ for a review). The PING model has been applied and validated in animals and recently in humans (pharmaco-MEG on using tiagabine and applying dynamic causal modelling) ^7^. We have previously demonstrated that sedation with propofol (as a representative drug with primarily GABA-ergic action) results in increased gamma band response (GBR), increased alpha power suppression, and a decrease in the amplitude of the stimulus-onset evoked response ^1^. This provided an insight into the possible separation of the neural generators of visual gamma oscillations ^8^ and the differential effects of propofol on those generating mechanisms. Propofol appeared to inhibit thalamo-cortical pathways resulting in decreased evoked^*^ visual responses while its intracortical GABAergic inhibition resulted in an enhanced induced^*^ gamma amplitude. This mechanistic discovery provides a potential biomarker to study and refine different pharmacological compounds that have similar clinical actions.

Dexmedetomidine produces sedation through mechanisms distinct from the commonly used GABAergic anaesthetic drugs (e.g. propofol and midazolam). Dexmedetomidine selectively acts on the α2-adrenergic receptors of the locus coeruleus, projecting to the preoptic area, which activates the inhibitory outputs to the arousal centres and results in sedation ^9^. Dexmedetomidine’s neurophysiological mechanisms, replicating ‘restorative sleep’ through activity on brainstem and normal sleep pathways, instead of the cortical suppression seen with GABAergic sedatives, may make it clinically advantageous especially in critically ill patients requiring long-term sedation ^10^. Easy arousability, with dexmedetomidine makes it particularly suitable for neurosurgical procedures (awake craniotomies) ^11^. Functional MRI studies to explain some of these clinical differences have indicated different effects on thalamo-cortical and cortico-cortical functional connectivity patterns ^12^. Dexmedetomidine also attenuates thalamic and cortical oscillations in the 30-200 Hz frequency bands ^13^. This effect on thalamic oscillations is less prominent with dexmedetomidine as compared to propofol, at frequencies > 50 Hz. While dexmedetomidine affects thalamic and cortical oscillations to a similar extent, propofol has a much greater effect on thalamic oscillations than cortical oscillations ^14^.

In this experiment, we used MEG to characterise and compare the effects of propofol and dexmedetomidine on visual cortical oscillations in a placebo-controlled, cross-over, single-blind study. Based on the current understanding of dexmedetomidine’s actions, i.e., primarily at the locus coeruleus leading on to the suppression of the cortex, we expected it to produce suppression of thalamocortical responses to the visual stimulus, with similar suppression of cortical activity. This would be different to propofol which is likely to produce a marked suppression of thalamocortical activity and also a marked (direct) inhibition of cortical activity due to its direct activity at widespread GABA receptors in those regions. We, therefore, hypothesised that unlike propofol, dexmedetomidine will cause a reduction in visual induced GBR, while propofol causes an increased induced GBR.

In addition, we aimed to characterise the effects of propofol and dexmedetomidine on motor cortical oscillatory activity during a simple finger abduction task. Previous work ^15-17^ on GABA-ergic activity on motor oscillations has been inconclusive. We hypothesised that motor cortex gamma activity generators would behave similarly to those of the visual cortex and contrasting effects of dexmedetomidine and propofol would be demonstrable.

## Methods

### Participants

Sixteen right-handed healthy male participants (mean age 27.3 years (SD 5.2, range 21-40) were recruited following a detailed screening procedure. The study was approved by Cardiff University’s Research Ethics Committee and all participants gave informed written consent. Medical screening was performed to ensure that all participants were in good physical and mental health and not on any regular medication (American Society of Anesthesiologists physical status 1). Any volunteer with complaints of regular heartburn or hiatus hernia, known or suspected allergies to propofol or dexmedetomidine (or its constituents), regular smokers, those who snored frequently or excessively, or who had a potentially difficult-to-manage airway were excluded.

### Monitoring, Drug Administration and Sedation Assessment

Throughout the experiments, all participants were monitored, as per anaesthetic standards, by two anaesthetists of which one was solely involved in monitoring. Participants were instructed to follow standard pre-anaesthetic fasting guidelines. Participants received either placebo (normal saline infusion), propofol or dexmedetomidine infusion in a pseudo-randomised design. These sessions were conducted over three separate visits, with each session separated from the next by a minimum of 72 hours to ensure complete clearance of the drug. For the control (normal saline) session, data were recorded starting 10 minutes into the infusion. Sedation level was assessed by the second anaesthetist (NS), using the modified Observer’s assessment of alertness/sedation scale (OAA/S) ^18^. Sedation endpoint was an OAA/S level of 4 (slurred speech with lethargic response to verbal commands).

### Propofol administration

Propofol (Propofol-Lipuro 1%, Braun Ltd., Germany) was administered using an Asena® - PK infusion pump (Alaris Medical, UK) using a target controlled infusion based on the Marsh-pharmacokinetic model as described in our previous work ^1^. While participants lay supine in the magnetically shielded room, infusion was started targeting an effect-site concentration of 0.6 mcg/ml. Once the target was reached, two minutes were allowed to ensure reliable equilibration. Drug infusion was then increased in 0.2 mcg/ml increments until the desired level of sedation was achieved.

### Dexmedetomidine administration

Dexmedetomidine (Dexdor®, Orion Corporation, Finland) was administered using a Graseby 3500® infusion pump (Smiths Medical, UK) controlled by a personal computer using the STANPUMP software using the Dyck pharmacokinetic model ^19^. Infusion was started targeting an effect site plasma concentration of 0.1 nanograms/ ml. Once the target was reached, five minutes were allowed to ensure further equilibration. Drug infusion was then increased in 0.1 nanograms/ ml increments until the desired level of sedation (OAA/S of 4) was achieved.

### Stimulation Paradigm

Once steady state sedation was achieved, participants were presented with a visual stimulus consisting of a vertical, stationary, maximum contrast, three cycles per degree, square-wave grating presented on a mean luminance background. Of a total 150 trials, 75 were displayed at maximum contrast, while the remaining 75 were displayed at 70% contrast. The radius of the grating was 8 degrees of visual angle, with a continually displayed, small, central, red fixation square. The grating patch was displayed for between 1.5 and 2s with 3s inter-stimulus interval (displaying a fixation square only). The stimulus was presented on a projection screen controlled by Presentation®. Stimuli were displayed by a Sanyo XP41 LCD back-projection system displaying at 1024 × 768 at 60 Hz. Participants were instructed to fixate on the red square throughout the trial and perform a finger abduction at grating-offset. Activity of the first dorsal interosseous muscle during finger abduction was recorded by both a bipolar EMG electrode placed either end of the muscle, and actual finger movement recorded by an optical displacement meter ^20^. Each recording session took, approximately, 15 minutes and was carried out before and during sedation.

### MRI acquisition

All participants had a structural MRI scan either as part of the study, or as participants in previous studies in Cardiff University Brain Research Imaging Centre (CUBRIC). Scans were conducted on a GE HDx 3T MR scanner with 8 channel head coil and followed a fast spoiled gradient echo (FSPGR) sequence with 1 mm isotropic voxel resolution. Co-registration with MEG data was achieved by matching fiducial coil positions recorded in MEG to the same location on MR images.

### MEG Acquisition and Analysis

Whole head MEG recordings were made using a CTF 275-channel axial gradiometer system (VSM MedTech) sampled at 1200 Hz (0–300 Hz bandpass). An additional 29 reference channels were recorded for noise cancellation purposes and the primary sensors analysed as synthetic third-order gradiometers ^21^. Three of the 275 channels were turned off due to excessive sensor noise. At the onset of each stimulus presentation a TTL pulse was sent to the MEG system. Participants were fitted with three electromagnetic head coils (nasion and bilateral pre-auriculars), which were localised relative to the MEG system immediately before and after the recording session. These were used for MRI/ MEG co-registration as described above.

### MEG Pre-Processing

Dataset markers were placed at the initiation of finger abduction, based on a shift in the amplitude of the optical displacement meter by three standard deviations above mean noise ^22^. Where noise masked a shift corresponding to a displacement, the EMG trace from the first dorsal interosseus was used. For the visual response, data were epoched into 4 s trials (from 2 s before to 2 s after the visual stimulus onset) to create a dataset containing only visual grating trials. For the motor response, data were epoched into 4.5 s trials consisting of 1.5 s pre- and 3 s post-finger abduction onset to create a dataset of only motor responses. Trials from both datasets were visually inspected for gross artifacts (head movements and muscle artifacts affecting a large number of sensors) and these trials were removed.

### Visual response source localisation

Visual stimulation, as used in this experiment, produces a typical response morphology (Fig 1-top panel): there is an initial transient broadband (50 to 100 ms) amplitude increase in the gamma frequency (40+ Hz) range (evoked response) followed by a longer-lasting elevation of gamma frequency amplitude in a narrower frequency range (induced response) ^23^.

**Figure 1:**
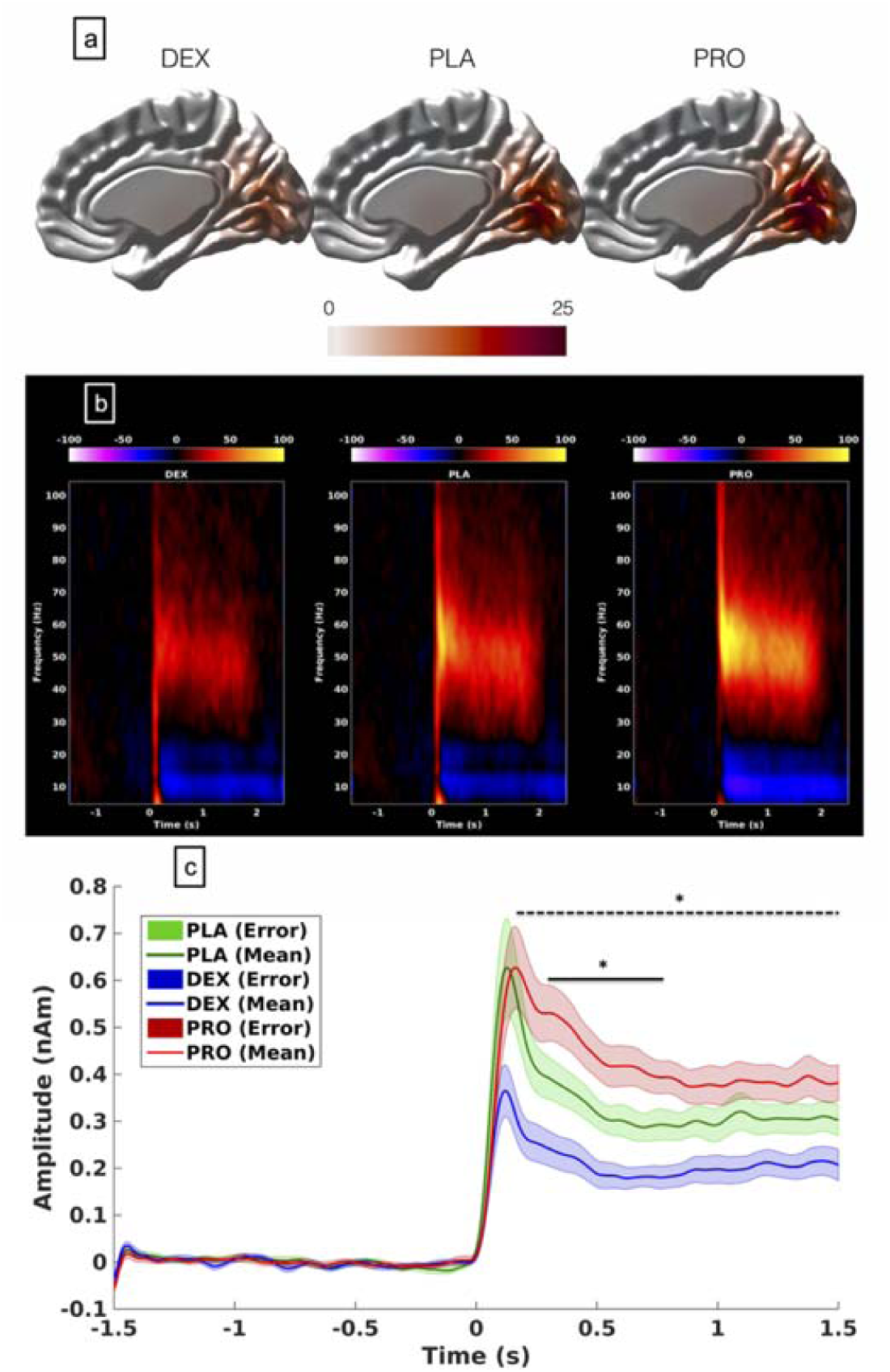
**a):** Grand-averaged source localisation of gamma oscillations (40–80 Hz) for awake and sedated states. Units are t statistics. PLA = placebo, DEX = dexmedetomidine, PRO = propofol. **b):** Grand-averaged time-frequency spectrograms showing source-level oscillatory amplitude (evoked + induced) changes following visual stimulation with a maximum contrast (100%) grating patch (stimulus onset at time = 0) during awake and sedated states. Spectrograms are displayed as percentage change from the pre-stimulus baseline and were computed for frequencies from 5 up to 150 Hz but truncated here to 100 Hz for visualisation purposes. **c):** Envelopes of oscillatory amplitude for the gamma (40–80 Hz). Time-periods with significant differences between the three conditions are indicated with a black bar (*p<0.05, shaded areas represent SEM). Colour: Blue-dexmedetomidine; Green-placebo; Red-propofol: Dotted bar-difference between dexmedetomidine and placebo; Bold bar-difference between propofol and placebo.

Two source localisations were performed on each dataset using synthetic aperture magnetometry, one for induced responses (SAM_ind_), and one for evoked responses (SAM_erf_) ^24^. Correspondingly, two global covariance matrices were calculated for each dataset, one for SAM_ind_ (40–80 Hz) and one for SAM_erf_ (0–100 Hz). Based on these covariance matrices, using the beamformer algorithm ^25^, two sets of beamformer weights were computed for the entire brain at 4 mm isotropic voxel resolution. A local-spheres ^26^ volume conductor model was derived by fitting spheres to the brain surface extracted by FSL’s Brain Extraction Tool ^27^.

For gamma-band SAM_ind_ imaging, virtual sensors were constructed for each beamformer voxel and student’s -t images of source power changes computed using a baseline period of - 1.5 to 0 s and an active period of 0 to 1.5 s. Within these images, the voxel with the strongest power increase (in the contralateral occipital lobe) was located. To reveal the time–frequency response at this peak location, the virtual sensor was repeatedly band-pass filtered between 1 and 150 Hz at 0.5 Hz frequency step intervals using an 8 Hz bandpass, 3rd order Butterworth filter ^23, 28^. The Hilbert transform was used to obtain the amplitude envelope and spectra were computed as a percentage change from the mean pre-stimulus amplitude (−1.5 to 0 s) for each frequency band. From these spectra, the time courses of alpha (8–15 Hz) and gamma (40–80 Hz) were extracted and submitted to non-parametric permutation tests using 5000 permutations and omnibus correction for multiple corrections ^29^. To examine pre-stimulus amplitudes the time-frequency spectra were recomputed with no baseline correction and the average amplitudes of alpha (8–15 Hz), beta (15–40 Hz) and gamma (40–80 Hz) in the pre-stimulus period (−1.5 to 0 s) were calculated.

For SAM_erf_, the computed evoked response was passed through the 0–100 Hz beamformer weights and SAM_erf_ images ^30^ were generated at 0.01 s intervals from 0.05 to 0.15 s. The image (usually 0.08 to 0.09 s or 0.09 to 0.1 s) with the maximal response in visual cortex was identified and the maximal voxel selected as the peak location for virtual sensor analysis. For time-domain analysis, the evoked field was computed for this virtual sensor (−0.2 to 0 s baseline, 40 Hz low-pass filter) and the peak amplitude and latency of the M100 and M150 responses were quantified. We also performed a spectral analysis of the evoked field using the same time-frequency techniques as above. The evoked frequency response in the 0 to 0.2 s period was obtained for each condition and analysed using the same statistical methodology.

### Motor response source localisation

Analysis of motor responses was procedurally similar to visual responses, except for the following differences. The motor paradigm elicits a narrow-band response between 60 and 90 Hz, ^31^ termed movement-related gamma synchrony (MRGS). This paradigm also elicits a robust bilateral beta de-synchronisation (movement related beta de-synchronisation: MRBD) followed by a beta-rebound (post-movement beta synchronisation/ rebound; PMBR), more prominent in the contralateral hemisphere. The beamforming and virtual sensor reconstruction procedure was repeated for each of these components with the beta range defined as 15–30 Hz. Guided by previous reports ^32^ the MRBD component was identified between 0.3 and 3 s while the PMBR component was identified between 1 and 2.5 s post finger-abduction. Virtual sensors were created separately for each participant and each condition (pre and post, for placebo, propofol and dexmedetomidine). As per the visual analysis, time frequency content was reconstructed at the virtual sensor location with the maximal relative response.

For further statistical analyses, a 3×2 repeated-measures ANOVA was used, with condition and time as factors (condition = dexmedetomidine, propofol or placebo, time = before or during-infusion), with the interaction term of primary interest. Paired t-tests were used for post-hoc between-group analyses. Results were corrected for multiple comparisons using Bonferroni’s correction. These are presented as ‘corrected’ in the subsequent text.

## Results

### Sedation level / dose

All participants were sedated to the desired level of mild sedation (OAA/S of 4). The mean plasma concentration of propofol required was 0.83 mcg ml^-1^ (SD 0.2 mcg ml^-1^) and for dexmedetomidine was 0.25 ng ml^-1^ (SD 0.12 ng ml^-1^). Both drugs reduced systolic BP (p < 0.005; Table 1) but had no effect on the heart rate. There was no difference in recorded head movement between the groups.

**Table 1.** Haemodynamic changes during infusions. There was a significant decrease in systolic blood pressure (SBP) in the propofol and dexmedetomidine groups (p < 0.005) but not in diastolic blood pressure (DBP) or heart rate (HR). Head movements between all groups-there were no differences. SD= Standard deviation. * p < 0.005

|  | **Placebo** |  | **Propofol** |  | **Dexmedetomidine** |  |
| --- | --- | --- | --- | --- | --- | --- |
|  | **Pre-**<br>**infusion** | **During-**<br>**infusion** | **Pre-**<br>**infusion** | **During-**<br>**infusion** | **Pre-**<br>**infusion** | **During-**<br>**infusion** |
| **SBP : Mean**<br>**(SD); mm Hg** | 123 (17) | 122 (11) | 125 (9) | 115 (13) * | 126 (11) | 118 (13) * |
| **DBP: Mean**<br>**(SD); mm Hg** | 70 (8) | 69 (10) | 70 (7) | 64 (10) | 70 (9) | 66 (7) |
| **HR: Mean**<br>**(SD); bpm** | 63 (7) | 63 (7) | 66 (9) | 64 (5) | 62 (9) | 60 (6) |
| **Head**<br>**movement:**<br>**Mean (SD); mm** |  | 1.57 (0.8) |  | 2.61 (3.3) |  | 3.29 (2.9) |

### Visual responses

The visual grating stimulus utilised here robustly elicits induced gamma responses in V1. The grand-averaged peak locations of the responses were located in adjacent source reconstruction voxels (4 mm voxel size) (Fig. 1a). This analysis found similar results from both maximum (100%) and low (70%) contrast grating patches and therefore only the results from the maximum contrast gratings are presented here. Data from low contrast gratings is presented in the Supplementary material (Fig. S1)

Figure 1b shows the group visual responses (representative subject datasets are presented in Supplementary data: Fig. S2). The virtual sensor reconstruction demonstrated changes in pre-stimulus gamma power between groups (F (2,30) = 3.17; p = 0.0035) (Supplementary data: Fig. S3) and therefore a ‘relative change’ (percentage from mean baseline) approach was utilised for analysis. In Figure 1c the extracted gamma (40–80 Hz) time-courses are plotted. For the high contrast stimulus, propofol resulted in a 44% increase in gamma amplitude, as compared to placebo, between 0.3 – 0.8 sec following the stimulus (t = 2.73, p = 0.027, corrected) while dexmedetomidine resulted in a 40% decrease in amplitude between 0.1 – 1.5 sec following the stimulus (t = −4.59, p = 0.004, corrected) (Fig. 1c). There was no change in peak-induced gamma frequency (F (2,30) = 0.074; p = 0.93) (Supplementary data: Fig. S4). Propofol (with high contrast gratings) resulted in an increased stimulus-induced alpha suppression by about 50% (t= 2.95, p = 0.02, corrected), however there was no change in alpha suppression by dexmedetomidine (Supplementary data: Fig. S5). There was no change in alpha suppression with either drug at low contrast settings (F (2,30) = 0.169, p = 0.173).

Figure 2 presents the time-averaged evoked responses. There were significant reductions in both the amplitude of the M100 (mean change 27%) (t = 6.9, p < 0.001, corrected) and M150 (mean change 52%) (t = −3.0, p = 0.018, corrected) components during propofol sedation. However, there were no differences between placebo and dexmedetomidine (M100: t = 1.19, p = 0.25; M150: t = −0.89, p = 0.388). We also noted significant slowing of the M100 component with both propofol (t = −4.2, p < 0.001, corrected) and dexmedetomidine (t = −4.6, p < 0.001, corrected). There was however no difference between the latencies of the M150 component (Supplementary data: Fig. S6).

**Figure 2:**
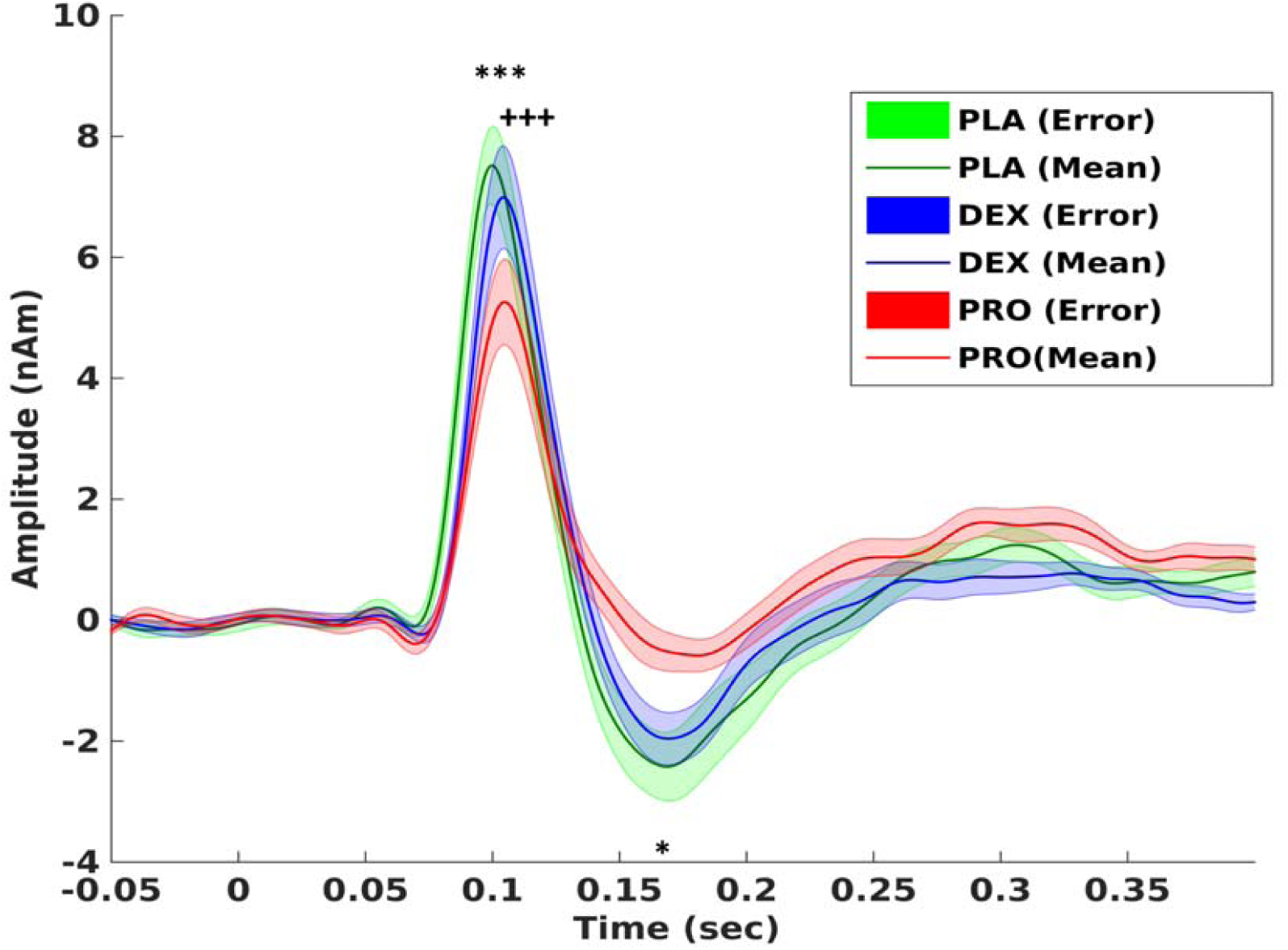
Visual responses: Source-level time-averaged evoked responses for placebo, propofol and dexmedetomidine. Significant differences were seen in M100 amplitudes: between propofol and placebo (***p<0.001); M100 latencies: between propofol and placebo (^+++^p<0.001) and between dexmedetomidine and placebo (^+++^p<0.001); M150 amplitudes: between propofol and placebo (*p<0.05). There were no significant changes between placebo and drugs on M150 latency. (2 tailed paired t-test, Bonferroni’s correction applied). Bold line represents means, shaded areas represent SEM.

The gamma band findings within the evoked response are presented in the supplementary material. Gamma band amplitude was reduced by 53% with dexmedetomidine (t = −3.58, p = 0.004, corrected) but not with propofol (t = 0.38, p = 0.7) (Supplementary data: Fig. S7a). There were no significant changes in the peak frequency, with the drugs (Supplementary data: Fig. S7b).

### Motor responses

The finger abduction task robustly elicits 3 components: a contralateral movement-related gamma synchrony, bilateral beta desynchrony followed by a contralateral beta-rebound (Figure 3a). Time–frequency analysis revealed an effect of the drugs on the baseline amplitude of the beta-rebound sensor (PMBR), MRGS and MRBD, hence subsequent time– frequency analysis of these sensors utilised a relative change approach. There were no significant changes with both drugs on the gamma amplitude or the MRBD, either the ipsilateral (right (BD_R_)) or contralateral (left (BD_L_)) sides (Fig. 3b). However, PMBR revealed increased power (92%) of ipsilateral (right (BR_R_)) beta rebound with dexmedetomidine (between 16 – 18.5 Hz, t = 2.6, p = 0.044, corrected) (Fig. 3c) but not on the contralateral side (left (BR_L_)). There was a non-statistically significant reduction (70%) with propofol in the contralateral (left (BR_L_)) PMBR (between 20 – 20.5 Hz) (t = −2.16, p = 0.05, uncorrected), but no change on the ipsilateral (right (BR_R_)) side.

**Figure 3:**
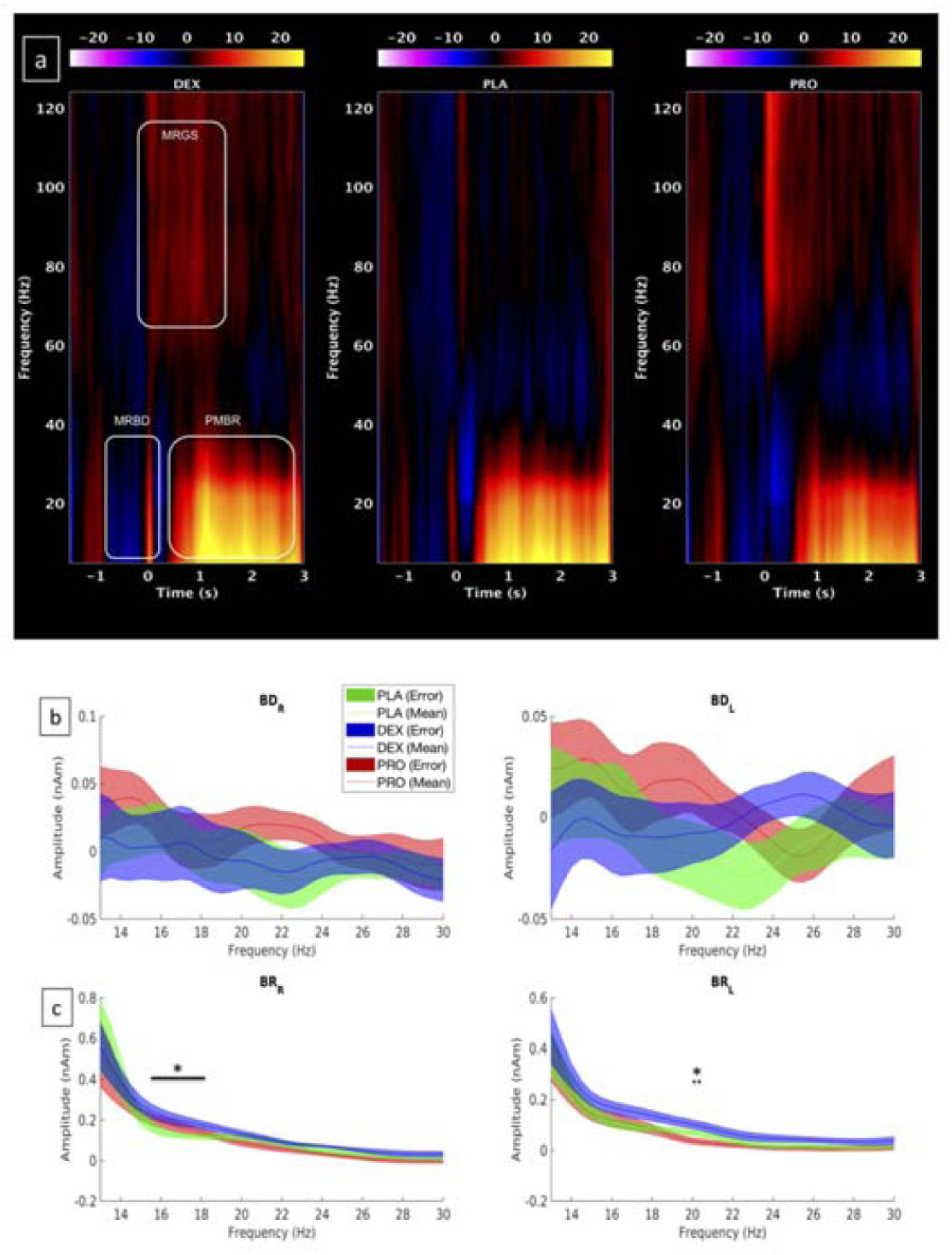
**<u>a)</u>** Time–frequency reconstructions for each condition (averaged over subjects) from M1 contralateral gamma-located virtual sensor, with amplitude depicted by heat-map colours. **<u>b)</u>** Movement related beta desynchronization (BD): Right (BD-R) and left (BD-L) sensors. **<u>c)</u>** Post movement beta rebound (BR): Right (BR-R) and left (BR-L) sensors. DEX = dexmedetomidine, PLA = placebo, PRO = Propofol. Colour: Blue-dexmedetomidine; Green-Placebo; Red-propofol: Bold bar-difference between dexmedetomidine and placebo. Dotted bar-difference between propofol and placebo. (*p<0.05).

## Discussion

This study reports the findings from a combined visuomotor MEG paradigm recorded during a placebo-controlled, crossover, single-blind study of the sedative effects of propofol (GABAergic sedative) and dexmedetomidine (non GABAergic sedative), on human cortical oscillations. The key findings were an increase in the stimulus-induced gamma band power with propofol, while there was a decrease in stimulus-induced gamma band power with dexmedetomidine. While propofol decreased the amplitudes of M100 and M150 evoked responses, dexmedetomidine had no effect on those. Dexmedetomidine, however, slowed the M100 response (increased latency) while propofol had no effect on M100 latency. Dexmedetomidine reduced stimulus-onset-evoked gamma band power, while propofol did not. In response to motor stimulation, neither drug had any significant effect on the motor gamma band power or the MRBD. However, dexmedetomidine increased PMBR power while propofol showed a tendency to decrease PMBR.

### Visual oscillatory differences

Our previous results ^1^, demonstrating the *in-vivo* modifiability of human GBR, were reproduced here using a more robust version of the stimulation protocol ^32^. The dissociation between the evoked and induced responses by propofol were considered akin to the dissociation between evoked responses and induced GBR representing separate thalamocortical and intracortical mechanisms, respectively, in the generation of high frequency oscillations ^8^; a finding which has also been demonstrated in human intra-cortical recordings ^33^. According to the PING model, GABAergic inhibition of interneuronal control may facilitate hypersynchronicity presenting as an increased power of the induced GBR. Suppression of both evoked responses and induced GBR by dexmedetomidine confirms a suppressive action on thalamocortical generators with no local intracortical facilitation, as hypothesised. These findings, in humans, provide further evidence of the differential roles of dexmedetomidine and propofol on thalamic and cortical oscillations.

Plourde and Arseneau ^13^, demonstrated that dexmedetomidine produced a dose-dependent attenuation of thalamic and cortical oscillations in the 30-200 Hz frequency bands. Dexmedetomidine attenuates both thalamic and cortical oscillations to a similar degree while propofol has a greater effect on the thalamic oscillations than cortical oscillations ^14^. EEG studies of the resting spectra have also revealed significant differences between dexmedetomidine and propofol. During moderate sedation, both dexmedetomidine and propofol increased spindle power; however, dexmedetomidine decreases global alpha, beta and gamma power, whereas propofol decreases alpha power in the occipital area and increases global beta and gamma power ^34^. In our experiment dexmedetomidine reduced the gamma power in the pre-stimulus baseline, while propofol did not result in any change (Supplementary data: Fig. 1) as would have been predicted based on previous literature ^34^. The novelty of our experiment is the demonstration of differences in the task-induced oscillatory changes, with visual-induced GBR being discriminatory between the representative GABA-ergic and non-GABA-ergic sedatives.

Alpha band activity is closely related to the gamma band activity, especially in the occipital cortex ^35^. While alpha activity is associated with an inhibitory function, in response to a task, it is suppressed to allow high frequency oscillations to transmit information. Thalamo-cortical neurons may be responsible for the generation and maintenance of the alpha-band oscillations ^36^. Modelling studies have suggested that the action of propofol, on these neurones, at unconsciousness producing doses, causes a suppression of posterior alpha and emergence of frontal alpha rhythms ^37^. Neural modelling of the changes in the resting EEG spectra during propofol anaesthesia suggest that these are caused by increased inhibition within local interneuron circuits ^38, 39^. An increase in alpha suppression with propofol replicates our previous findings ^1^, reflecting increased local GABA-ergic inhibitory effects. This increased alpha suppression was not seen with our low contrast visual task. Luminance contrast has been shown to linearly increase the gamma band response and leave beta (13-30 Hz) band response unaffected ^40^. It is unclear if alpha suppression is inversely related to luminance contrast although there were no differences found between high and low contrast in the placebo group. Our results, however, suggest that propofol induced alpha suppression enhancement may be related to the contrast of the visual stimulus. Dexemdetomidine, does not alter local excitatory-inhibitory balance and therefore demonstrated no effect on task-induced alpha suppression.

Reduction in M100 (amplitude and latency) and M150 (amplitude) with propofol were similar to those reported in our previous work ^1^. Dexmedetomidine did not decerease the amplitudes of either M100 or M150, although it increased the latency of M100. Dexmedetomdine has been shown not to affect evoked repsonses (including visual), although most studies have been performed during intra-operative use (under anaesthesia with other drugs) ^41^. We were unable to find any data on dexmdetomidine’s independent effects on visual evoked responses. Our results therefore suggest that at the doses studied in this experiment, dexmedetomdine’s effect on evoked resposnes are substanially less than that of propofol and it only slows the M100 evoked responses without altering its amplitude or the M150 response.

### Motor oscillatory differences

The results of the motor task related oscillations have also revealed differences between the two sedatives. Finger movement (or similar motor activity) produces a decrease in amplitude of the ongoing beta oscillations in the primary motor cortex (MRBD). Following the termination of the movement, an increase in beta oscillatory power above the pre-movement oscillatory amplitude (PMBR) is observed ^42^. In addition to these beta band changes, there is a transient increase in power in the gamma frequency range, specifically in the 60–90 Hz frequency, temporally coincident with movement onset ^22^. GABAergic mechanisms have been proposed to explain these findings, both for the gamma ^43^ and beta ^44, 45^ band oscillations. This follows the PING model, whereby these oscillations are facilitated by increasing the inhibitory drive of GABAergic interneurons, via GABA-A receptors. As the inhibitory post synaptic potential (IPSP) decay-time is lengthened, this reduces the frequency of the locally oscillating neuronal network population. Consequently, this serves to facilitate the recruitment of principal cells to the oscillating population, giving rise to an increase in the amplitude of the oscillatory power, as the participating neuronal pool is increased. Increases in GABA-A receptor driven beta frequency oscillatory power occur in M1 layers III and V ^44^ the predominant generators of signals measured using EEG and MEG approaches.

We had predicted an increase in motor gamma activity with propofol, similar to the increase seen with visual gamma. However, there were no changes in the motor gamma activity with either propofol or dexmedetomidine. Interestingly, a similar lack of changes in motor gamma with GABA modulators (diazepam ^16^ and tiagabine (GABA transporter inhibitor, which increases synaptic GABA levels) ^17^) has been reported. Motor gamma was increased by ketamine ^46^ and alcohol (both GABA and glutamatergic activity)^15^ suggesting that glutamatergic influence may be the dominating influence on these oscillations. Our findings therefore reinforce the importance of glutamatergic rather than GABAergic effects, within the excitation-inhibition model influencing the modulation of motor cortical gamma oscillations.

MRBD is considered a non-specific state of movement preparation, starting before movement, from the contralateral M1 and then becoming bilateral ^47^. MRBD, on both contralateral and ipsilateral sides, was unaffected by either propofol or dexmedetomidine. Previously reported increases in MRBD with diazepam ^16^, and tiagabine ^17^ but not with propofol, suggests that propofol did not increase GABA-A activity to the extent required for this effect to be detectable. Unlike, MRBD, the proposed significance of PMBR includes a sensory re-afference to motor cortex following movement ^48^, stabilising current motor output and, therefore, in preventing initiation of new movements ^49^ and reflecting neural processes that evaluate motor error in the context of the prior history of errors ^50^. Propofol showed a tendency to reduce PMBR (contralateral) while dexmedetomidine increased PMBR activity (ipsilateral). Interestingly, contrary to other motor beta findings, it has been suggested that PMBR may be a non-GABA-A mediated effect, as evident by absence of effect with diazepam ^16^. Indeed a decrease with tiagabine ^17^ and propofol (in this experiment)(which have some GABA-B agonist activities ^51^), suggests that this may be a marker of enhanced GABA-B activity. PMBR activity tends to be localised to the contralateral side and the significance on the ipsilateral rebound is less clear. Motor-related tasks which do not involve actual movement such as motor imagery ^52^ have shown preferential ipsilateral beta synchrony, while reading ^53^ and foot movement planning ^54^ have shown beta synchrony over both ipsilateral and contralateral M1 areas. Such ipsilateral synchrony may be interpreted as a correlate of a deactivated or actively inhibited motor area neurons wherein enhanced inhibition of the ipsilateral motor area occurs via the transcollosal fibre system ^55^. An increased ipsilateral PMBR with dexmedetomidine, may reflect a more rapid re-afferentation process, which in turn may be a factor in the rapid arousal, as seen clinically, with dexmedetomidine.

We conclude that dexmedetomidine and propofol affect the visual and motor cortical oscillations differently. At equi-sedative doses, propofol increases visual stimulus-induced GBR while dexmedetomidine decreases it. Propofol reduces M100 and M150 amplitudes while dexmedetomidine has no effect on these evoked response amplitudes. Both drugs increased the latencies of M100 but not the M150. Dexmedetomidine reduced stimulus-onset-evoked gamma band power, while propofol did not. PMBR power is increased by dexmedetomidine while it is reduced by propofol. The findings of this experiment provide a mechanistic link between the known receptor physiology of these sedative drugs and the differences in their clinical effects. Better understanding of the neurophysiologic correlates of sedation, based on receptor physiology is likely to help understand the different components of consciousness better, help develop more reliable monitoring tools and help develop anaesthetic drugs with a better safety profile.

## Details of authors contributions

N.S.: study conception, study design, recruitment of volunteers, study conduct, data collection, data analysis, interpretation of results, writing the first draft of the manuscript and manuscript revision.

A.D.S.: data analysis, interpretation of results, writing the first draft of the manuscript and manuscript revision.

A.B.: recruitment of volunteers, study conduct, data collection.

L.R.: recruitment of volunteers, study conduct, data collection.

K.D.S.: study design, data analysis, interpretation of results and manuscript revision.

J.E.H.: study design and manuscript revision.

R.G.W.: study design, interpretation of results and manuscript revision.

S.D.M.: study design, recruitment of volunteers, study conduct, data collection, data analysis, interpretation of results, writing the first draft of the manuscript and manuscript revision.

## Acknowledgements

None

## Footnotes

* Evoked vs Induced responses Evoked (also known as ‘spike’) responses represent neural activation that occurs at the same time, phase-locked with respect to stimulus or task onset (or offset) from trial to trial. Induced (also known as ‘sustained’) responses are time-dependent variation of the amplitude of oscillations within a frequency band of interest. These may reveal effects that occur systematically across trials but are less strictly time-locked and are not phase-locked. Therefore, they typically disappear in the time-domain averaging commonly employed to analyse evoked responses.

