## Supplementary for "A comparison of GABA-ergic (propofol) and non-GABA-ergic (dexmedetomidine) sedation on visual and motor cortical oscillations, using magnetoencephalography"

**Supplementary Material**

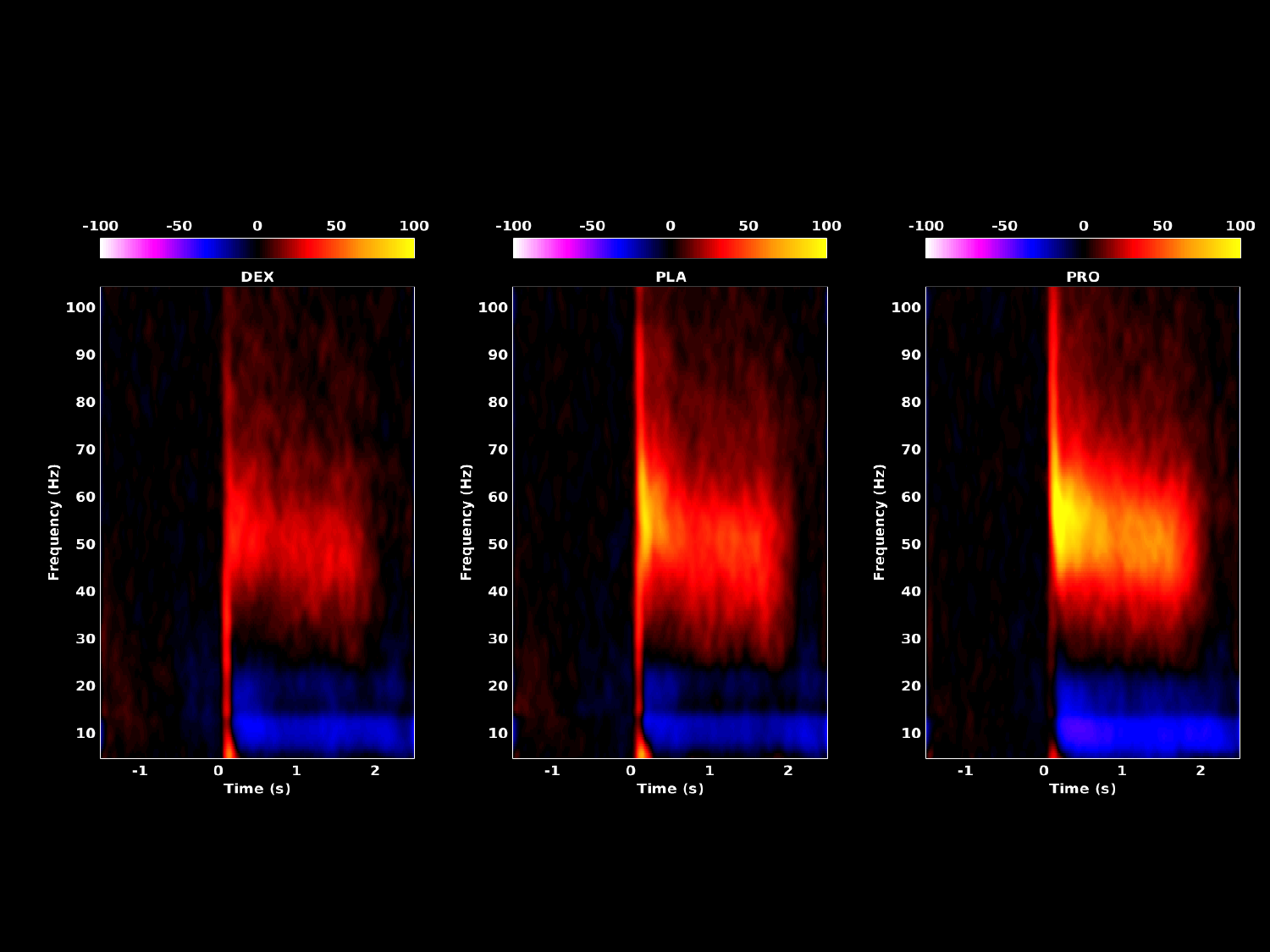

Figure S1: Grand-averaged time-frequency spectrograms showing source-level oscillatory amplitude (evoked + induced) changes following visual stimulation with a 70% contrast (low contrast) grating patch (stimulus onset at time = 0) during awake and sedated states. Spectrograms are displayed as percentage change from the pre-stimulus baseline and were computed for frequencies from 5 up to 150 Hz but truncated here to 100 Hz for visualisation purposes. DEX = dexmedetomidine, PLA = placebo, PRO = Propofol.

Results were similar to those obtained with maximum (100%) contrast gratings in the gamma frequency band. In the alpha frequency band there were no group differences.

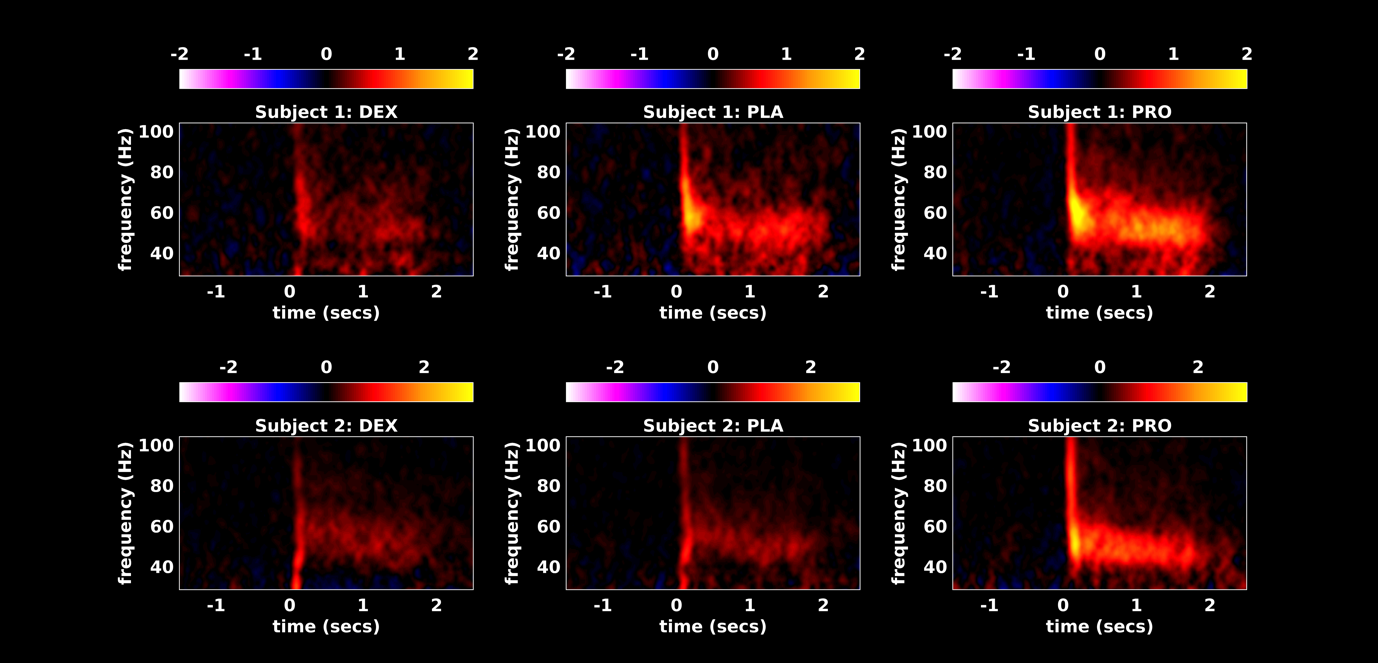

Figure S2: Representative time-frequency spectrograms showing source-level oscillatory amplitude (evoked + induced) changes, in the gamma band, following visual stimulation with a 100% contrast grating patch (stimulus onset at time = 0) during awake and sedated states. Spectrograms are displayed as mean changes from the pre-stimulus baseline DEX = dexmedetomidine, PLA = placebo, PRO = Propofol.

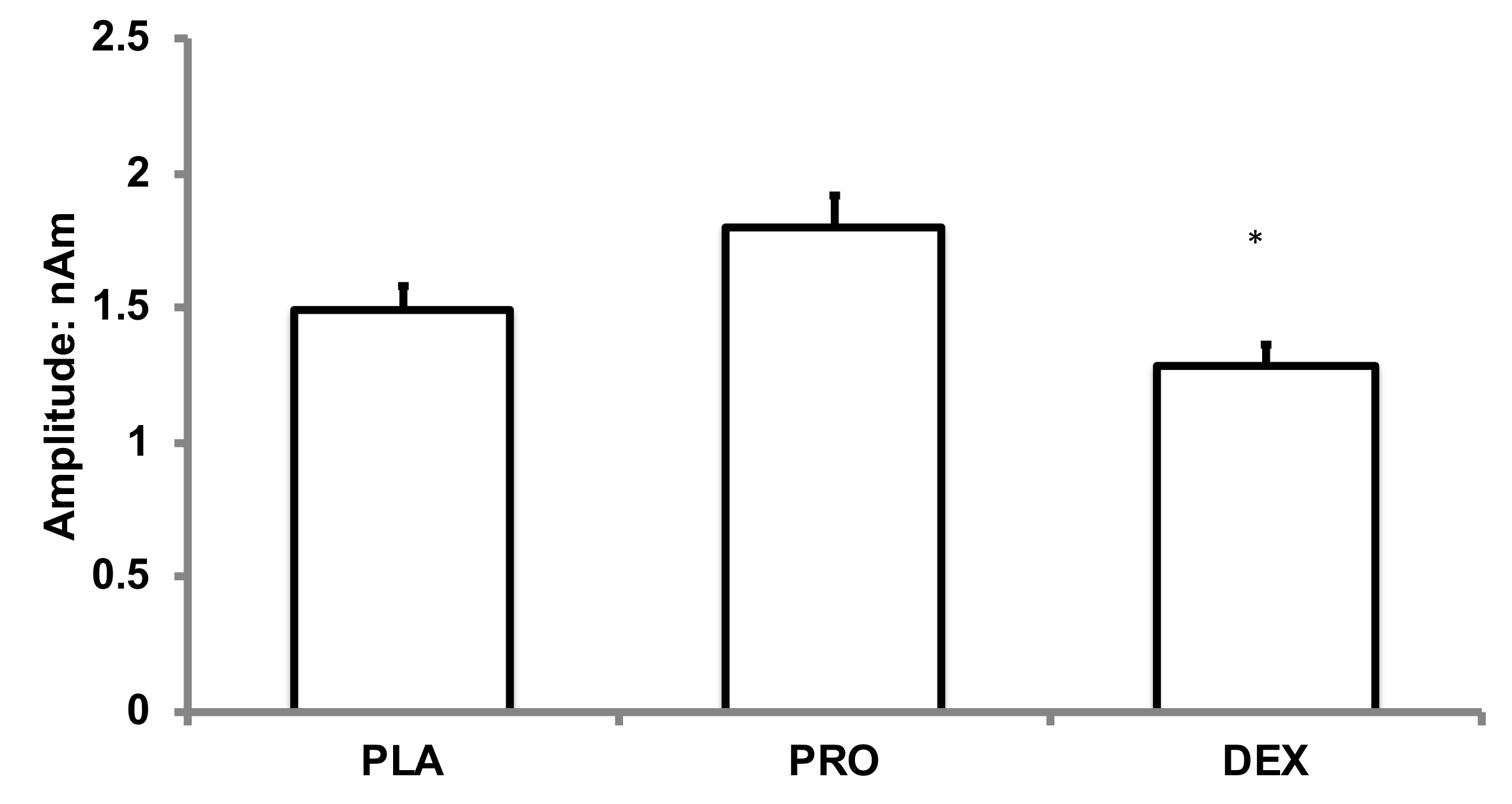

Figure S3: Bar charts showing baseline (pre-stimulus) gamma band amplitude changes (high contrast). PLA = placebo, DEX = dexmedetomidine, PRO = propofol. *p<0.05, compared to placebo; error bars represent SEM.

Figure S4: Bar charts showing peak frequency in the induced- gamma band (high contrast) PLA = placebo, DEX = dexmedetomidine, PRO = propofol. Error bars represent SEM. There were no significant differences.

Figure S5: Bar charts showing changes in stimulus induced- alpha band (8-13 Hz) power (change in suppression) during high contrast visual task. PLA = placebo, DEX = dexmedetomidine, PRO = propofol. *p<0.05, compared to placebo; bars represent SEM. There were no significant differences during the low contrast visual task.

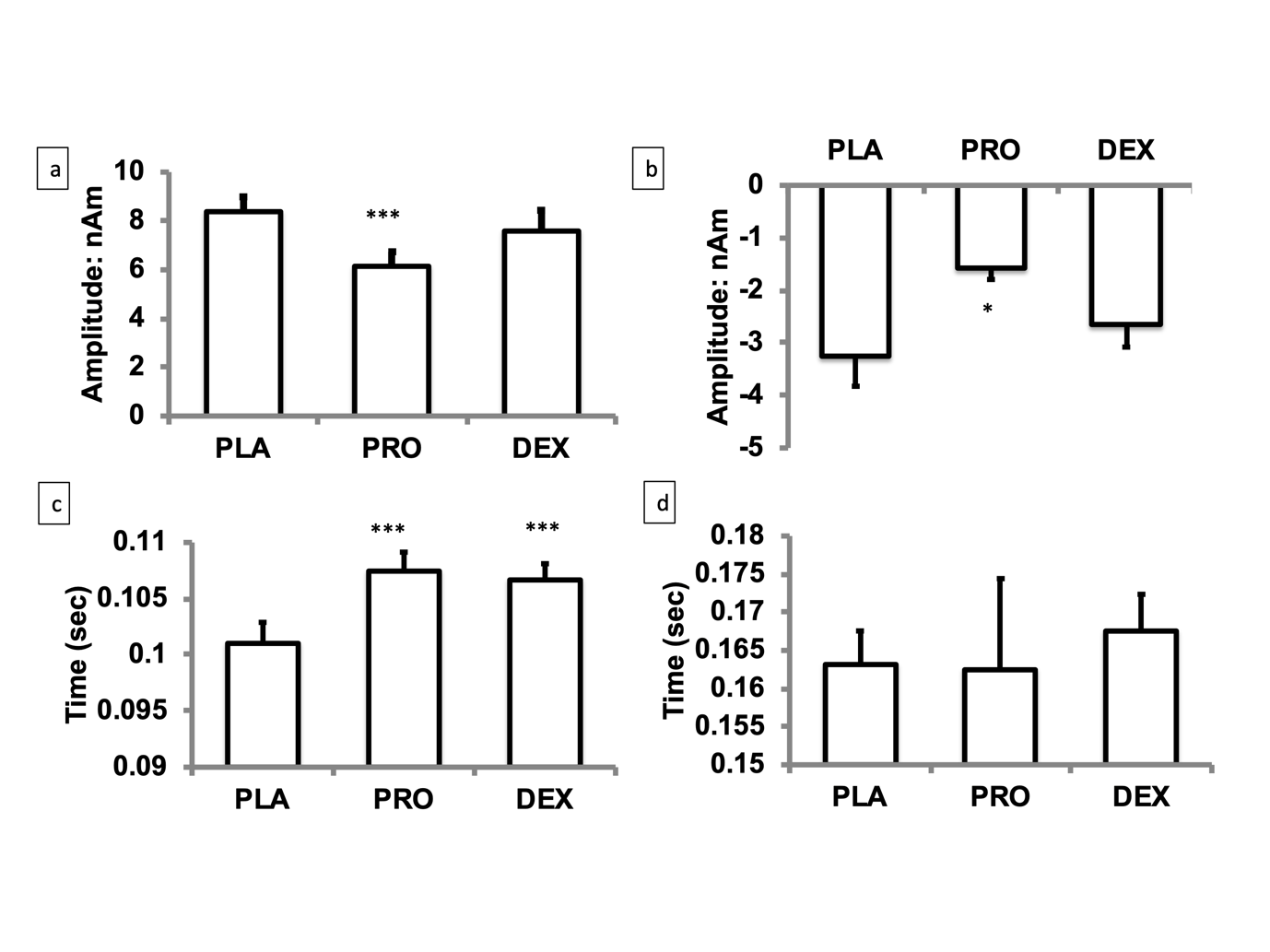

**Figure S6:** Bar charts showing source-level time-averaged evoked responses for placebo, propofol and dexmedetomidine. PLA = placebo, DEX = dexmedetomidine, PRO = propofol. Significant differences were seen in (a) M100 amplitudes (b) M100 latency and (c) M150 amplitudes (2 tailed paired t-test; difference between drug and placebo: *p<0.05, ***p<0.001, bars represent SEM). No differences were seen in (d) M150 latency.

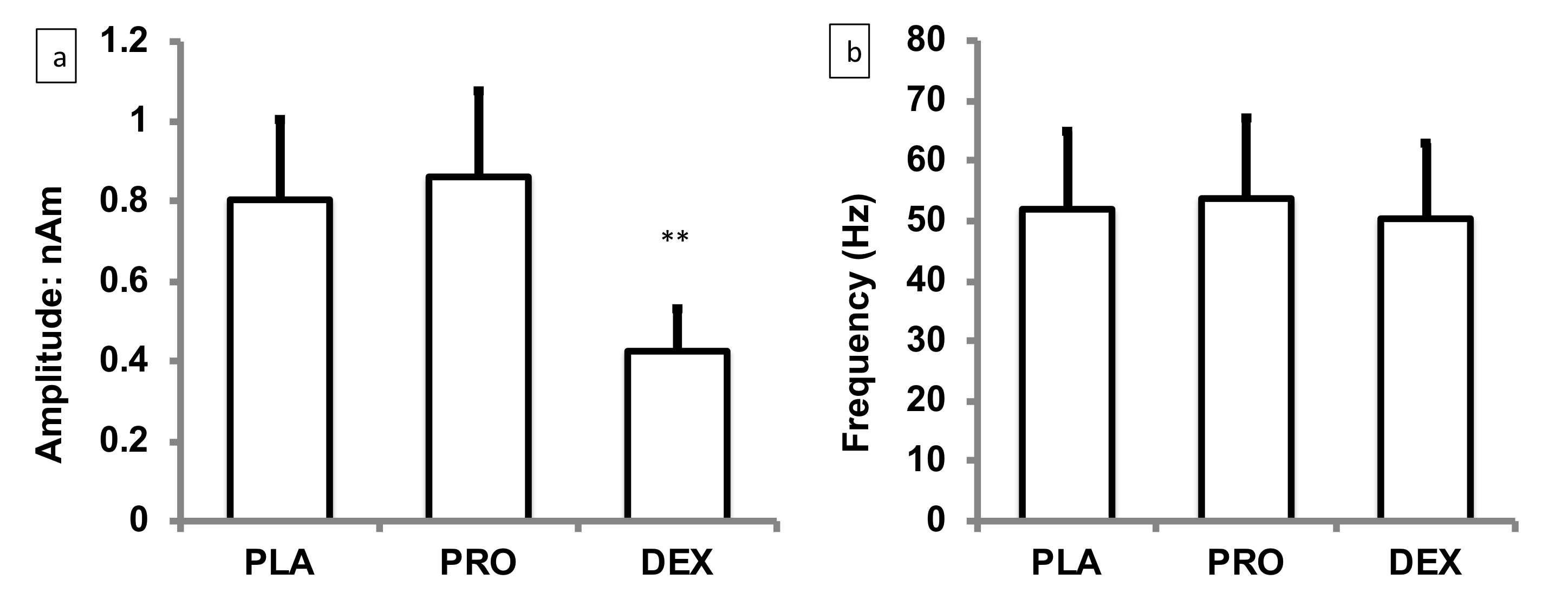

Figure S7: Visual responses during the evoked phase of the high contrast visual task, in the gamma band range. PLA = placebo, DEX = dexmedetomidine, PRO = propofol. a) amplitude of gamma power , b) peak frequency. **p<0.005, compared to placebo; bars represent SEM.
